# Automated pupillometry to detect command following in neurological patients: A proof-of-concept study

**DOI:** 10.1101/525212

**Authors:** Alexandra Vassilieva, Markus Harboe Olsen, Costanza Peinkhofer, Gitte Moos Knudsen, Daniel Kondziella

**Author notes:** **Corresponding author:** Daniel Kondziella, MD, MSc, dr.philos., FEBN, Department of Neurology, Rigshospitalet, Copenhagen University Hospital, Blegdamsvej 9, DK-2100 Copenhagen. **Ethics approval:** The Ethics Committee of the Capital Region of Denmark approved the study (journal-nr.:H-18045266).

## Abstract

**Abstract:** *Background:* Levels of consciousness in patients with acute and chronic brain injury are notoriously underestimated. Paradigms based on electroencephalography (EEG) and functional magnetic resonance imaging (fMRI) may detect covert consciousness in unresponsive patients but are subject to logistical challenges and the need for advanced statistical analysis.

*Methods:* To assess the feasibility of automated pupillometry for the detection of command following, we enrolled 20 healthy volunteers and 48 patients with a wide range of neurological disorders, including 7 patients in the intensive care unit (ICU), who were asked to engage in mental arithmetic.

*Results:* Fourteen of 20 (70%) healthy volunteers and 17 of 43 (39.5%) neurological patients, including 1 in the ICU, fulfilled prespecified criteria for command following by showing pupillary dilations during ≥4 of 5 arithmetic tasks. None of the 5 sedated and unconscious ICU patients passed this threshold.

*Conclusions:* Automated pupillometry combined with mental arithmetic appears to be a promising paradigm for the detection of covert consciousness in people with brain injury. We plan to build on this study by focusing on non-communicating ICU patients in whom the level of consciousness is unknown. If some of these patients show reproducible pupillary dilation during mental arithmetic, this would suggest that the present paradigm can reveal conscious awareness in unresponsive patients in whom standard investigations have failed to detect signs of covert consciousness.

## Introduction

It can be difficult to assess if patients with acute brain injury are conscious by means of standard clinical examinations alone because these patients must be sufficiently aroused and able to mobilize motor function (Schnakers et al., 2009b,a; Laureys et al., 2010; Di et al., 2014; Kondziella et al., 2016). Thus, standard neurological assessment often misclassifies unresponsive patients as being in a vegetative state (VS, aka. unresponsive wakefulness syndrome, UWS) (Schnakers et al., 2009b). This has important implications for prognosis and puts these patients at risk of unjustified withdrawal of life-sustaining therapy (Demertzi et al., 2011; Turgeon et al., 2011; Ong, Dhand & Diringer, 2016; Harvey et al., 2018).

Our limited knowledge of disorders of consciousness contributes to this dilemma. It is still not widely recognized that many patients are entirely unable to interact with their environment because of complete motor paralysis, despite being minimally conscious (minimal conscious state, MCS) or even fully conscious (Kondziella et al., 2016). This state of covert consciousness in completely paralyzed patients has been termed cognitive motor dissociation (Schiff, 2015). Owen and co-workers were the first to document cognitive motor dissociation in a landmark paper from 2006 (Owen et al., 2006). Herein, the authors showed that a young traffic accident victim without any signs of consciousness at the bedside, thereby fulfilling clinical criteria of VS/UWS, was able to follow commands simply by modulating her brain’s metabolic activity as measured by functional magnetic resonance imaging (fMRI) (Owen et al., 2006). Thus, in the past 15 years consciousness paradigms based on fMRI and electroencephalography (EEG) that circumvent the need for motor function have been developed (Monti et al., 2010; Cruse et al., 2011; King et al., 2013; Sitt et al., 2014; Stender et al., 2014; Rohaut et al., 2017; Vanhaudenhuyse et al., 2018). However, although these technologies may detect covert consciousness, fMRI- and EEG-based paradigms are labor-intensive, expensive, logistically challenging and not readily available in the intensive care unit (ICU) (Weijer et al., 2015; Kondziella et al., 2016). A cheap and fast, easy-to-interpret point-of-care test for consciousness assessment at the bedside is clearly needed.

Automated pupillometry may just be this test. The pupillary reflex is a polysynaptic brainstem reflex under cortical modulation, i.e. cognitive processes such as decision making and mental arithmetic produce pupillary dilation (Kahneman & Beatty, 1966; Marquart & de Winter, 2015). Hence, pupillary responses following mental arithmetic have been used to establish communication with patients with the locked-in syndrome and to detect command-following in 1 patient in MCS (Stoll et al., 2013). However, these were patients with chronic brain injury many months or years after the injury, and the technical equipment used was complex, involving a fixed bedside camera and a computer screen for the display of visual instructions (Stoll et al., 2013). Here, we wished to assess whether a convenient hand-held pupillometer and a simpler paradigm with vocal instructions allow reliable measurements of pupillary dilation during mental arithmetic as a sign of command following, and hence conscious awareness, in a wide range of neurological patients admitted for in-hospital care.

## Methods

### Objectives

We aimed to evaluate a paradigm for the assessment of conscious awareness in patients with neurological disorders admitted to in-patient hospital care. To this end, we assessed pupillary dilation following mental arithmetic in neurological patients in the ICU and neurological ward, as well as in healthy volunteers. Sedated unconscious patients served as negative controls. We hypothesized that most (but not necessarily all) healthy volunteers and conscious neurological patients would be able to cooperate and show pupillary dilation during mental arithmetic, whereas unconscious and sedated patients would not.

### Study population

We collected a convenience sample of 48 neurological patients admitted to the neuro-ICU and neurological wards at Rigshospitalet, Copenhagen University Hospital, including unsedated ICU patients with spontaneous eye opening in MCS or better (conscious state; CS). Five unconscious and deeply sedated/comatose patients in the ICU were recruited for negative control. Levels of consciousness were estimated following standard neurological bedside examination by a board-certified neurologist experienced in neurocritical care and according to established criteria (Giacino et al., 2018). Twenty healthy volunteers served as positive controls.

### Automated pupillometry and mental arithmetic paradigm

The integrity of the pupillary light reflex of both eyes was checked using the NPi^®^-200 Pupillometer (NeurOptics, Laguna Hills, CA 92653 USA). We documented the neurological pupil index (NPi), which is a proprietary pupillometry sum score from 0-5, with ≥3 indicating physiological limits (including a maximal difference between the 2 eyes of <0.7) (Chen et al., 2011; Larson & Behrends, 2015; Peinkhofer et al., 2018), pupillary diameters before and after light exposure, percentage change of pupillary diameters, and pupillary constriction and dilation velocities. Patients with non-physiological values were excluded. Then, we used the PLR^®^-3000 pupillometer (NeurOptics, Laguna Hills, CA 92653 U.S.A) to track pupillary size over time (approximately 3-5 minutes in total), while asking the patients to engage in mental arithmetic (**Figure 1**). Each patient was asked to calculate a series of 5 arithmetic problems of moderate difficulty (21 × 22, 33 × 32, 55 × 54, 43 × 44, 81 × 82; approx. 30 seconds each) with rest periods (30 seconds) in-between. A subgroup of patients was given arithmetic problems of lesser difficulty (4 × 46, 8 × 32, 3 × 67, 6 × 37, 7 × 43; approx. 15 seconds each, with 15 seconds rest periods). We carefully explained all participants that pupillary dilation is induced solely by the efforts associated with mental arithmetic, and that it was irrelevant for our study if their calculations were correct or not. Hence, participants were instructed not to reveal the results of their mental arithmetic but to pay attention to the task and make an honest effort.

**Figure 1.**
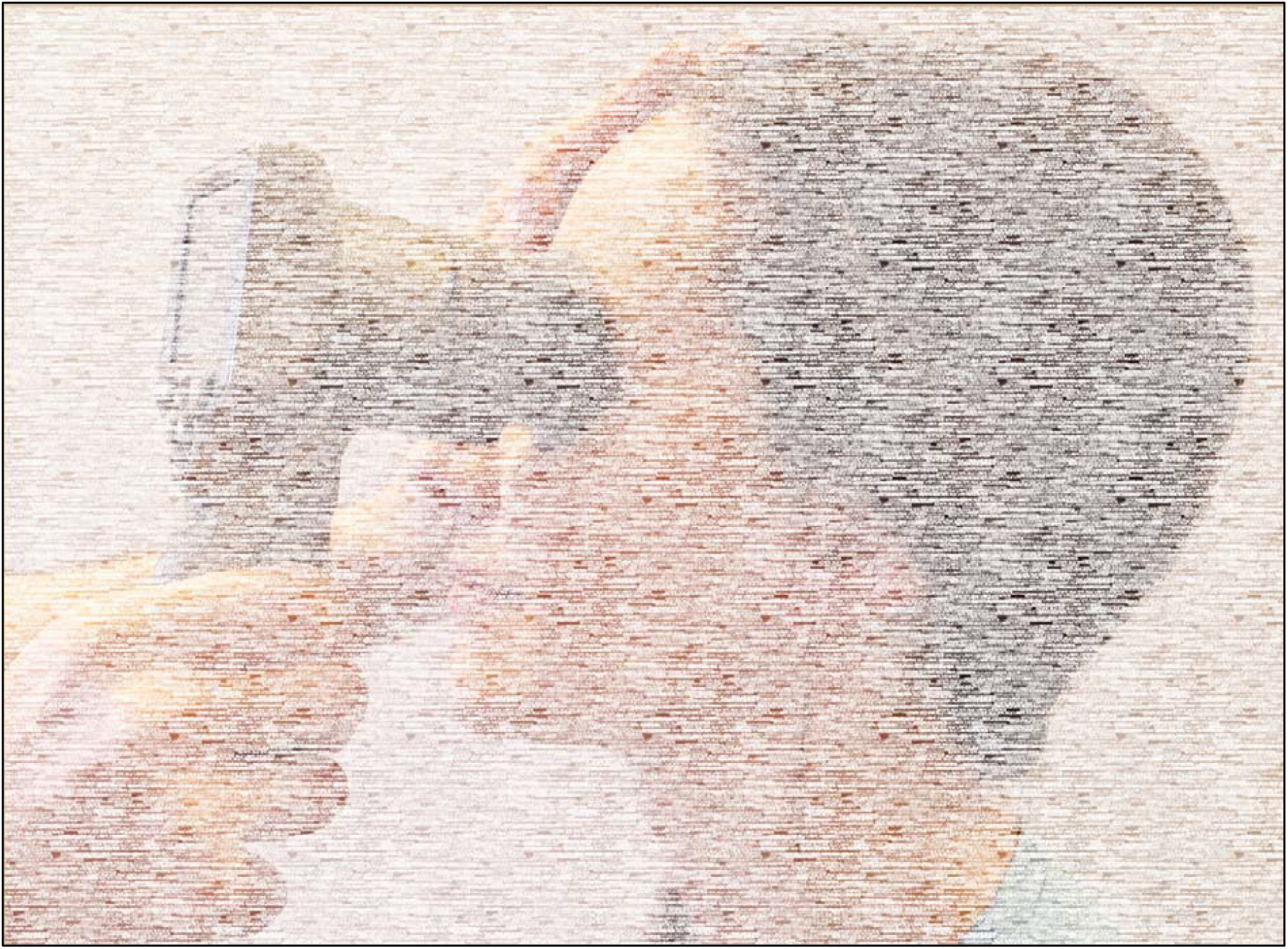
We used the PLR^®^-3000 pupillometer (NeurOptics, Laguna Hills, CA 92653 U.S.A), an automated handheld device, to track pupillary size, while asking patients and healthy volunteers (one shown here) to perform mental arithmetic.

### Outcome measures

Outcome measures included pupillary diameters during periods of mental arithmetic (intervention) and relaxation (rest periods).

### Statistical analysis

Data were analysed using R (R 3.4.1, R Development Core Team [2008], Vienna, Austria). Pupillary measurements were visually assessed for quality control in a run chart. Pupillary diameter changes in each of the five mental arithmetic tasks (intervention) were assessed by comparing the period of intervention with the periods of relaxation (rest periods) before and after. Successful pupillary dilation during intervention was defined as a significantly larger median pupillary size during mental arithmetic compared to the immediate rest periods prior and after (p-value < 0.01; Wilcoxon signed-rank test, followed by Bonferroni correction). We deemed command following to be successful when a participant showed pupillary dilation in at least four of the five mental arithmetic tasks (80%).

### Ethics

The Ethics Committee of the Capital Region of Denmark approved the study (journal-nr.:H-18045266). Written consent was obtained from all participants or their next-of-kin (unconscious or minimally conscious ICU patients). Data were anonymized and handled according to the European Union’s Data Protection Law. The pupillometry device used in the present study (NPi^®^-200 Pupillometer (NeurOptics, Laguna Hills, CA 92653 USA) was on loan from the manufacturer; however, neither the manufacturer, nor the vendor were involved in the design or conduct of the study, data analysis or writing of the manuscript, and the authors did not receive any other monetary or non-monetary benefits.

## Results

We enrolled 68 participants: 20 healthy controls, 41 neurological patients on the ward and 7 patients in the neuro-ICU. Diagnoses reflected a wide spectrum of neurological disorders, including cerebrovascular, neuromuscular, epilepsy, trauma, neuroinfections, and multiple sclerosis. Baseline pupillary function was normal in all participants and did not differ between healthy controls (mean NPi score 4.3 ± 0.39) and neurological patients (4.3 ± 0.32; p=0.812). **Table 1** shows demographic data.

**Table 1.** This tables shows demographic data, including neurological diagnoses. “Stroke” includes hemorrhagic and ischemic stroke, “neuromuscular” includes Guillain-Barré syndrome, chronic inflammatory demyelinating polyradiculopathy, Isaacs’ syndrome, Pompe’s disease and multifocal motor neuropathy, “epilepsy” denotes epilepsy with or without structural cause on magnetic resonance imaging. CS = conscious, ICU = intensive care unit, IQR = interquartile range, MCS = minimally conscious state, N = number of subjects, SAH = subarachnoid hemorrhage, TBI = traumatic brain injury. * All mental arithmetic tasks involved 2×2-ciffered calculations (e.g. 33 × 32), except for 1×2-ciffered calculations (e.g. 8 × 32) in neurological patients indicated with (*). ** This unsedated ICU patient in MCS with a pontine hemorrhagic stroke was examined twice at 7 days interval but failed to show command following during mental arithmetic in both sessions. *** Other diagnoses, not listed above, include relapsing remitting multiple sclerosis, unspecified sensory disturbances, brain abscess, anoxic-ischemic encephalopathy and hemangioblastoma

|  | Healthy volunteers | Neurological patients | Neurological patients * | ICU patients, MCS or CS | ICU patients, coma |
| --- | --- | --- | --- | --- | --- |
| N | 20 | 20 | 21 | 2 | 5 |
| Female | 10 (50%) | 9 (45%) | 8 (38%) | 2 (66%) | 2 (40%) |
| Age in years, median (IQR) | 34.5 (29-47) | 60.5 (51-68) | 50 (41-70) | 34 (34-34) | 62 (55-64) |
| Stroke | - | 2 | 2 | 1 ** | 0 |
| SAH | - | 0 | 1 | 0 | 2 |
| TBI | - | 0 | 0 | 0 | 2 |
| Epilepsy | - | 5 | 2 | 0 | 0 |
| Neuromuscular | - | 9 | 10 | 1 | 0 |
| Other *** | - | 4 | 6 | 0 | 1 |

Fourteen of 20 (70%) healthy volunteers fulfilled the prespecified criteria for successful command following, whereas this was the case for only 16 of 41 (39%) neurological patients on the ward, 1 of 2 unsedated patients in the ICU, and 0 of 5 comatose/sedated patients in the ICU. Healthy controls had higher rates of command following than neurological patients (risk ratio 1.81, 95% CI 1.13-2.99; z-statistic 2.48; p=0.013; excluding sedated ICU patients). Reducing the degree of difficulty of the mental arithmetic task did not change the proportion of neurological patients passing criteria for command following (8/20 patients, 40% vs. 8/21 patients, 38%).

Pupillary dilation was seen in 65 of 100 (65%) measurements in healthy controls; in 58 of 100 (58%), respectively, 72 of 105 (68.6%, simpler tasks) measurements in neurological patients on the ward; and in 7 of 15 (46.67%) measurements in unsedated ICU patients. By contrast, larger pupillary diameters were noted during 7 out of 25 (28%) measurements in comatose/sedated ICU patients (negative control group), consistent with chance occurrence.

**Table 2** provides details. Examples from healthy controls and neurological patients are given in **Figure 2** and **Figure 3.** Anonymized raw data are available in the *online supplemental files*.

**Table 2.** This table depicts the rate of successful command following by mental arithmetic in healthy volunteers, conscious neurological patients on the ward, minimally or fully conscious patients in the ICU, and comatose/sedated ICU patients. Successful command following was defined by ≥4 significant pupillary dilations during 5 mental arithmetic tasks. CS = conscious, ICU = intensive care unit, MCS = minimally conscious state, N = number of subjects. * All mental arithmetic tasks involved 2×2-ciffered calculations (e.g. 33 − 32), except for 1×2-ciffered calculations (e.g. 8 × 32) in neurological patients indicated with (*). ** This unsedated ICU patient in MCS with a pontine hemorrhagic stroke was examined twice at 7 days interval but failed to show command following during mental arithmetic in both sessions

|  | Healthy volunteers | Neurological patients | Neurological patients * | ICU patients, MCS or CS | ICU patients, coma/sedation |
| --- | --- | --- | --- | --- | --- |
| **N** | **20** | **20** | **21** | **2** | **5** |
| 0 significant | 2 | 3 | 1 | 1** | 1 |
| 1 significant | 2 | 1 | 1 | 0 | 2 |
| 2 significant | 0 | 2 | 1 | 1** | 1 |
| 3 significant | 2 | 6 | 10 | 0 | 1 |
| 4 significant | 13 | 5 | 4 | 0 | 0 |
| 5 significant | 1 | 3 | 4 | 1 | 0 |
| Successful | 70% (n=14/20) | 40% (n=8/20) | 38% (n=8/21) | 33% (n=1/3) | 0% (n=0/5) |

**Figure 2.**
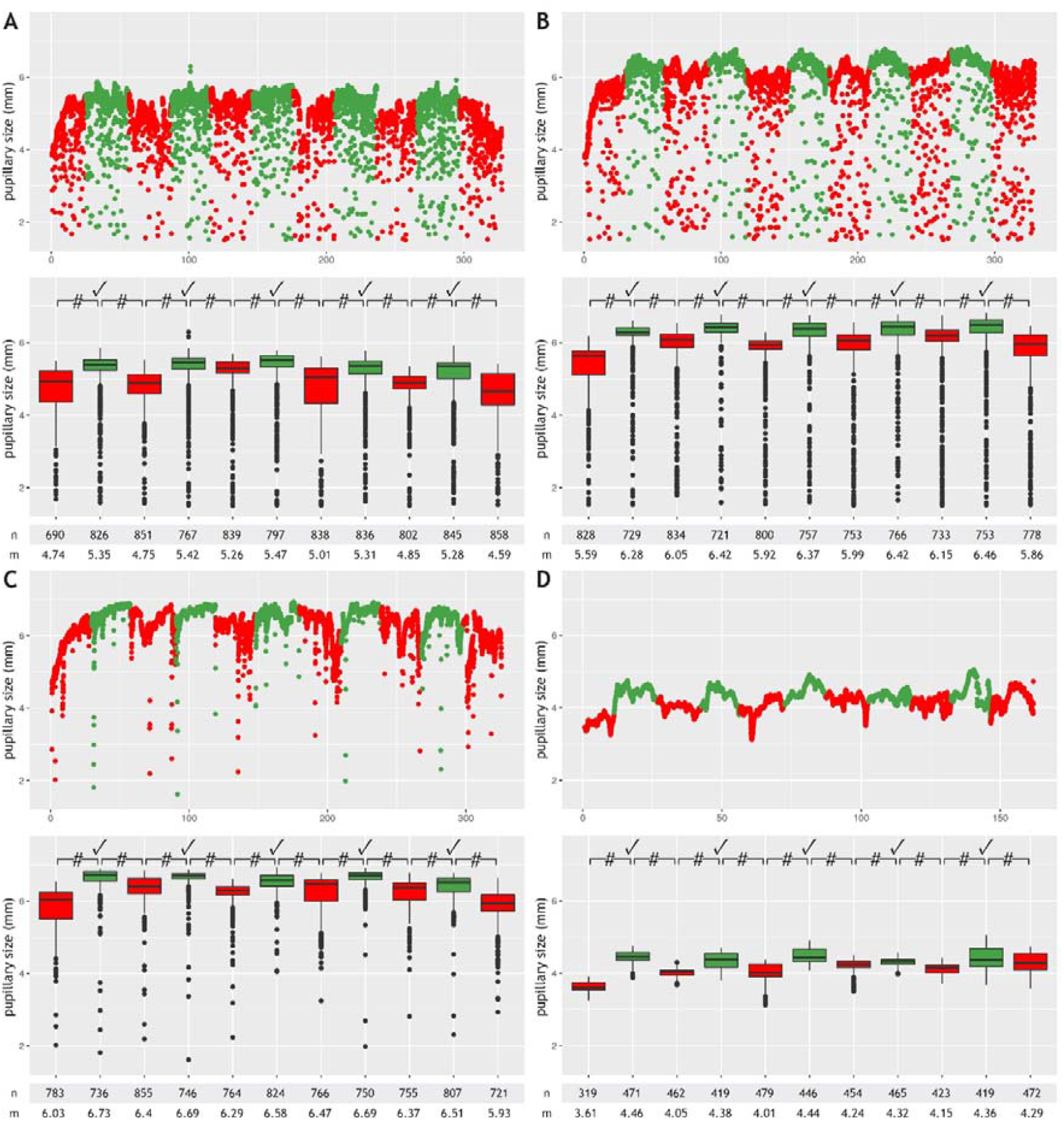
This figure shows results from participants with successful command following, detected by automated pupillometry, during a mental arithmetic paradigm: two patients admitted to the neurological ward with a diagnosis of multifocal motor neuropathy (**A**), respectively, Guillain-Barré syndrome (**B**), a healthy participant (**C**), and a conscious 34-year old male with the pharyngeal-cervical-brachial variant of Guillain-Barré syndrome admitted to the ICU (**D**). Minor artifacts due to blinking or eye movements are seen in A-C, but not D (probably because of facial and oculomotor nerve palsies). Periods with mental arithmetic are shown in green, rest periods in red. Pupillary sizes during mental arithmetic were significantly larger (p-value < 0.0001) than during rest periods, consistent with pupillary dilation, in all five tasks for each of the four participants. # = p-value < 0.0001; ⃠ = pupillary dilation; n = number of measurements; m = median pupillary size, mm = millimeter

**Figure 3.**
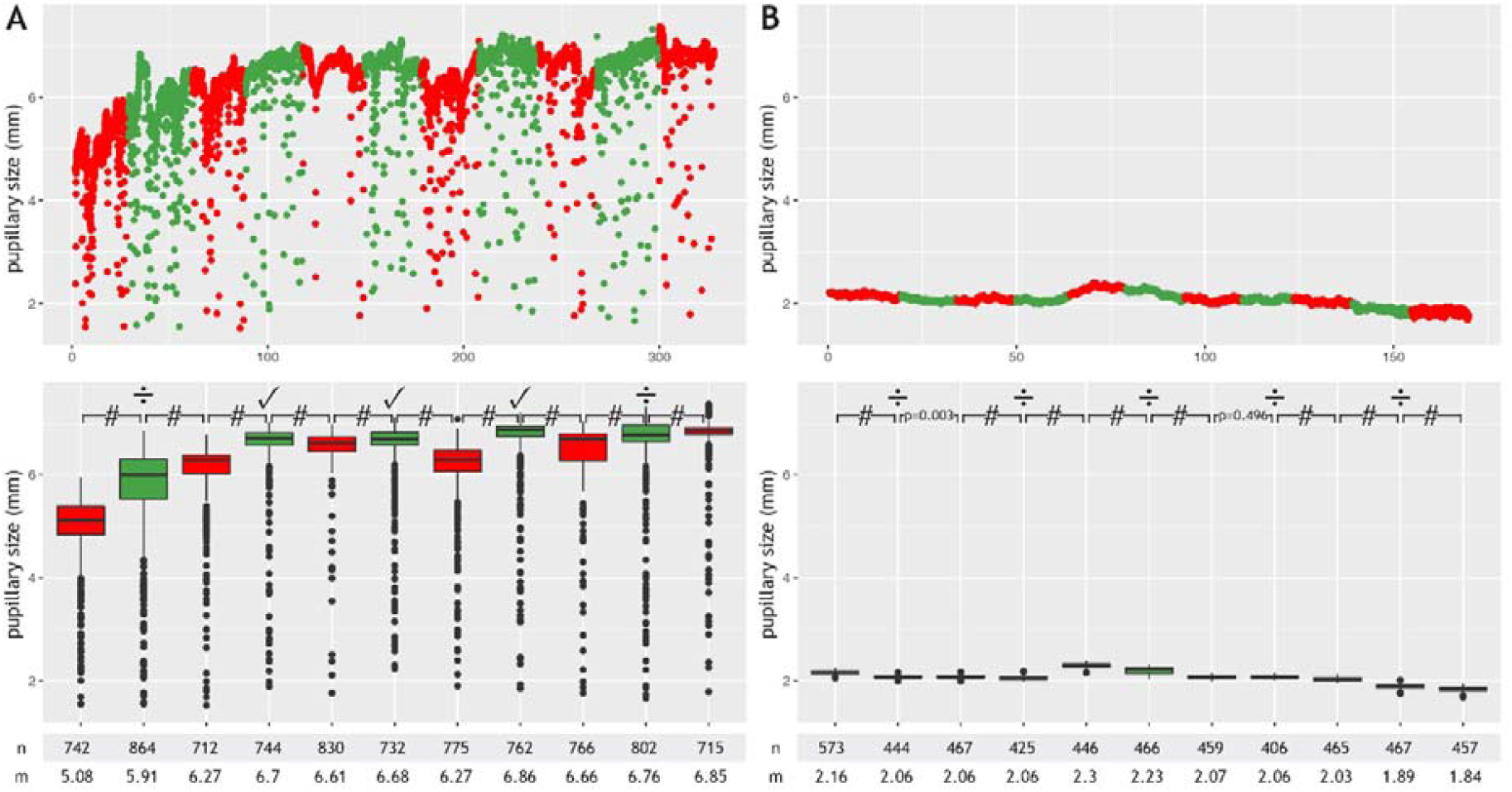
This figure depicts results from a healthy volunteer with unsuccessful command following (**A**) and a 62-year old male in the ICU with subarachnoid hemorrhage and deep sedation (Richmond Agitation-Sedation Scale score of −4) who served as a negative control (**B**). Minor artifacts due to blinking or eye movements are seen in A, but not B (because of sedation-induced impairment of the blink reflex). In the healthy volunteer, pupillary dilation was noted in only three out of five mental arithmetic tasks, which did not meet our prespecified criteria for successful command following (≥4 pupillary dilations, 80%). Minor random fluctuations in the pupillary diameter are seen in the unconscious sedated ICU patient. # = p-value < 0.0001; ⃠ = pupillary dilation; ÷ absence of pupillary dilation; n = number of measurements; m = median pupillary size, mm = millimeter

## Discussion

Cognitive and emotional processes evoke pupillary dilation in both humans and non-human primates, reflecting vigilance, arousal and attention (Laeng, Sirois & Gredebäck, 2012; Schneider M. et al., 2016; Becket Ebitz & Moore, 2017; McGarrigle et al., 2017; Foroughi, Sibley & Coyne, 2017). Hence, pupillary diameters may serve as an index of brain activity and mental efforts (or lack hereof). Here, we employed mental arithmetic as a paradigm for patients and healthy volunteers to control and maximize pupil dilation to signal command following. We found that a short session of mental arithmetic with simple verbal instructions, without prior training, revealed command following as detected by a handheld automated pupillometry device in 70% of healthy volunteers and 40% of conscious neurological patients.

Seventy % command following in healthy people may seem low, but this figure is very consistent with what has been reported with active EEG- and fMRI-based paradigms. For instance, in one study, 9 of 12 (75%) healthy controls produced EEG data that could be classified significantly above chance (Cruse et al., 2011); in another study, 12 of 16 healthy subjects had EEG responses to motor imagery [75.0% (95% CI: 47.6–92.7%)] (Edlow et al., 2017). Similarly, 11 of 16 healthy volunteers [68.8% (95% CI: 41.3–89.0%)] demonstrated responses within supplementary motor areas and premotor cortices when examined by a motor imagery fMRI paradigm (Edlow et al., 2017). Again, 7 of 10 (70%) healthy subjects were able to demonstrate covert command following in another motor imagery fMRI study (Bodien, Giacino & Edlow, 2017). Thus, many healthy people cannot cooperate in active paradigms. Of note, however, mental arithmetic seems to generate the most robust activation in the majority of healthy subjects for both EEG and fMRI (Harrison et al., 2017).

To assess the feasibility of mental arithmetic and pupillometry in the clinical setting, we pragmatically enrolled a large variety of patients with neurological disorders. Not surprisingly, the rate of command following was substantially lower - around 40% - either because of mild cognitive dysfunction related to the underlying neurological condition, the mental stress associated with being admitted to hospital, the higher median age, a lower level of education or a combination of these factors, although we did not examine this specifically. It is likely that allowing training sessions until patients feel they can cooperate might have yielded a higher success rate. Although we enrolled only two unsedated ICU patients, one of them successfully participated in our paradigm, which corroborates the feasibility of our approach. Also, as expected, none of the sedated ICU patients met our prespecified criteria of command following despite spontaneous fluctuations in pupillary diameters.

## Conclusions

Here we have shown that a fast and easy paradigm based on automated pupillometry and mental arithmetic is able to detect command following in healthy volunteers and conscious patients with neurological disorders admitted to in-hospital care. As a next step, we plan to focus on unsedated non-communicating ICU patients in whom the level of consciousness is unknown. Patients who show reliable pupillary dilation following mental arithmetic are likely to be conscious, whereas absence of pupillary dilation is a poor predictor of lack of consciousness. We suggest that our paradigm can be helpful to identify conscious awareness in non-communicating patients with acute brain injury for whom traditional bedside examination and laboratory investigations have failed to detect signs of covert consciousness. Compared to EEG and fMRI, pupillometry for the detection of command following, and ultimately covert awareness, would offer several advantages in the clinical setting, including quick and convenient assessment at the bedside, simple analysis and low costs.

Author contribution
AV, MHO, CP: acquisition of data, analysis and interpretation, critical revision for important intellectual content, approval of final manuscript; GMK: critical revision for important intellectual content, approval of final manuscript; DK: study concept, acquisition of data, analysis and interpretation, writing of the manuscript, critical revision for important intellectual content, approval of final manuscript

